# Prospects of raising baobab (*Adansonia digitata* L.) to fruiting in two years

**DOI:** 10.1101/2021.08.19.457013

**Authors:** Kenneth Fafa Egbadzor, Jones Akuaku

**Author notes:** Both authors contributed equally to the work.

## Abstract

Baobab is a very important plant with several uses, serving as food security crop and has the potential to raise income of farmers. However, the plant is undomesticated. We have the objective of domesticating the plant and promote its cultivation and utilization. Seeds were treated with sulphuric acid for early germination and grafted with matured scions suspected to be fruiting for several years. Twenty-two of the seedlings were transplanted on the field at Ho Technical University campus. Unfortunately, only four survived bushfire and destruction by stray animals. Surprisingly, one of the four surviving trees anthesis within twenty-eight months after seed treatment (twenty months after transplanting). This is the earliest ever reported anthesis of baobab. With better agronomic practices such as supplementary irrigation, insect pest control, manuring and protection from stray animals, baobab could start fruiting within two years of planting. This would alleviate fear of long maturity period and motivate farmers to go into baobab plantation. This would help diversify agriculture in many African countries, increase income and food security and contribute towards the attainment of the sustainable development goals of the United Nations.

## Introduction

Baobab (*Adasonia digitata* L.) domestication has been suggested by a number of authors in the past due to its importance (Jamnadass *et al*, 2011; Simbo *et al*, 2012; North *et al*, 2014). It is considered one of the most important African indigenous fruit trees needing domestication (Gebauer *et al*, 2016). This call started many years ago by Michel Adanson in whose honor the plant was scientifically named *Adansonia digitata* (www.powbab.com/pages/baobab-tree). There was a report on consortium of scientist working on baobab domestication in 2005 (Jensen *et al*, 2011). However, there are no results indicating sustained interest and work on the domestication of the plant at the moment.

In 2008, baobab was accepted as novel food in Europe and the United States of America (North *et al*, 2014). That might have triggered the high demand for its products in Europe and North America (Venter and Witkowski, 2010). Dependence on wild baobab products cannot meet world demand. Thus, domesticating the plant is imperative to sustainable supply to meet local and external demand. Revisiting the idea of baobab domestication is therefore, a step in the right direction.

Among the issues concerning baobab domestication and cultivation are the problem of seed dormancy and long juvenile period of the plant. Exposure to high degrees of heat in nature strongly influence germination of baobab seeds (Lautenschlager *et al*, 2020). Artificially, research has shown that treating the seed with concentrated sulphuric acid gives early and uniform germination. Boiling water and other treatments have also been reported to aid germination by Venter (2016) and El-Bably and Rashed (2018) among others authors.

In the case of long juvenile period, the use of vegetative planting material is most appropriate as known in other plants. Vegetative propagation methods thus need to be explored to reduce the maturity period of the plant to attract farmers to cultivate it. Vegetative propagation also results in producing true to type plants and will be particularly important in baobab to help manage wide diversity that could result from seed propagation (Agbohessou *et al*, 2020). The vegetative methods ever reported in baobab include grafting, cutting and *in vitro* propagation (Agbohessou *et al*, 2020).

Grafting has been the mostly used among the vegetative methods. Anjarwalla *et al* (2017) grafted scions on one to two-year-old rootstocks while Mbora *et al* (2008) used three months old rootstocks. Graft success was high for all ages of scions used. The most important consideration seems therefore to be the rootstock and scion sizes. Every available method has to be explored in working towards early fruiting of baobab in order that the long maturity period would not be discouraging factor in considering baobab farming. The objective of the study therefore, was to use sulphuric acid scarification for seed germination and grafting to investigate how early baobab can flower and bear fruit.

## Materials and Method

### Seed treatment

Baobab seeds were soaked in 95 % sulphuric acid (H_2_SO_4_) for six hours after which they were removed and washed thoroughly in tap water. Washed seeds were placed on wet paper towel and covered with same. Germination (protrusion of radicles) occurred five days after placing in the paper towel.

### Nursing and nursery practices

Germinating seeds were planted in soil contained in nursery bags (20 cm height and 15 cm diameter). Nursery bags were placed under citrus trees that allowed only partial penetration of sunlight. Watering was done whenever necessary. Pruning of the citrus tree was done after five weeks to allow more light to the baobab seedlings. There was no application of insecticides nor fertilizers at the nursery stage.

### Grafting

Scions were taken from matured baobab trees suspected to be fruiting for several years and grafted on the seedlings (rootstock) seven months after seed treatment. Top cleft method was used in grafting.

### Transplanting and other cares

Transplanting was done at Ho Technical University campus in October, 2019, that was three months after grafting. Apart from watering at the time of transplanting, no other irrigation was done apart from natural precipitation. Two hand-trowel full of poultry manure was supplied two months after transplanting. Broad spectrum insecticide was applied to control leaf eating insects. Weed was regularly controlled.

## Observations and Discussion

Baobab radicle emergence which was taken to be seed germination occurred five days (average) after treatment with sulphuric acid. All the pregerminated seeds transferred to soil filled nursery bags emerged from the soil. The seedlings grew rapidly and were grafted within seven months after planting with about 90 % success. Grafting at seven months is significant improvement over one to two years rootstock recommended by Anjarwalla *et al* (2017). However, this can be further reduced considering the three-month-old rootstocks reported by Mbora *et al* (2008). At three months, the rootstocks used in this trial were too small for the available scions. Different rootstocks can be explored to select faster growing ones to shorten the grafting time.

Growth was generally rapid during the raining seasons and very slow during the dry seasons characterized by dropping of leaves. First anthesis was observed 824 days (28 months or two years, four months) after seed treatment as shown in Table 1. The period from transplanting to anthesis was only 595 days (about 20 months). This observation in less than two years after transplanting is an extra ordinary. Jensen *et al* (2011), reported anthesis in baobab to be between eight and twenty-three (8-23) years when planted by seed or three to five (3-5) years when grafted. However, in recent publication, Agbohessou *et al* 2020 stated that result of grafted baobab to fruiting has not yet been documented. There are two similar but different issues here. One is flowering and the other fruiting. In both cases, our observation of flowering is earlier than ever reported. Prior to the current study, the earliest ever reported flowering of grafted baobab was by Sidibe and Williams (2002) and Mbora *et al* (2008). They reported that grafted baobab can flower after three years. Our observation is earlier than the three years, thus making it currently, the earliest baobab flowering ever reported.

**Table 1:**
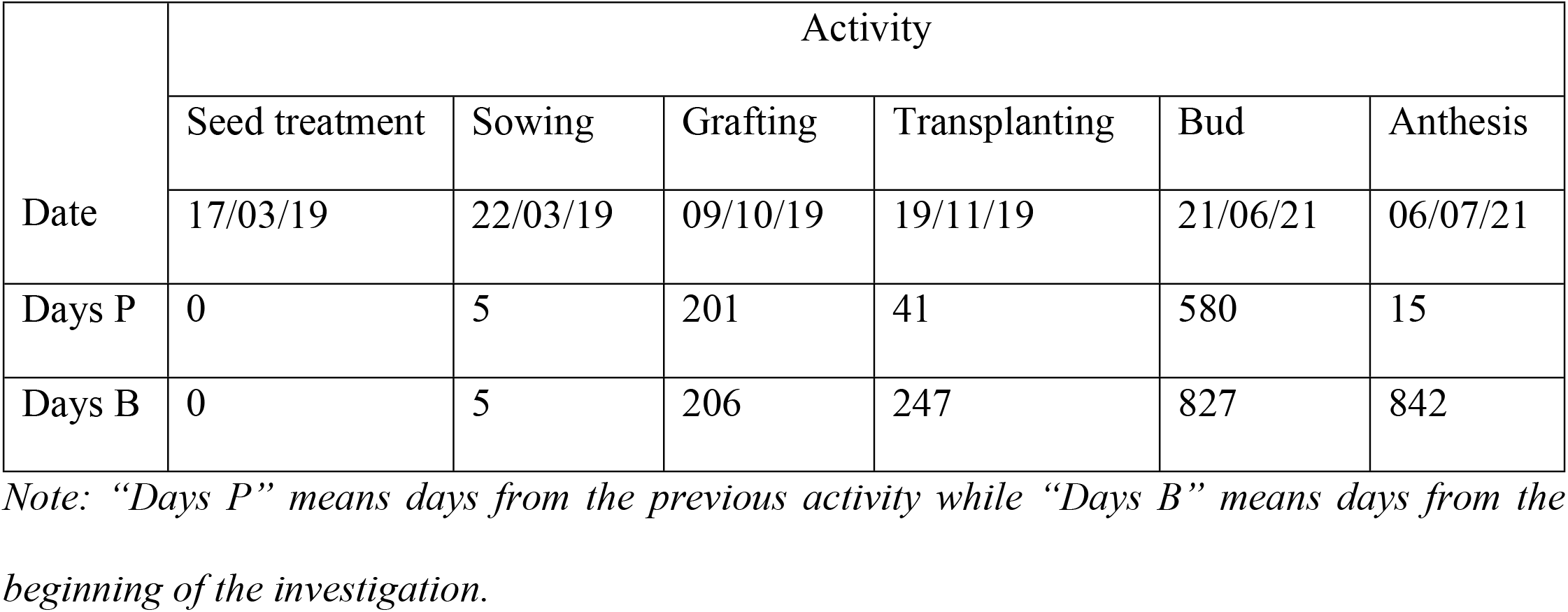
Timeline of activities and observations

| Date | Activity |  |  |  |  |  |
| --- | --- | --- | --- | --- | --- | --- |
|  | Seed treatment | Sowing | Grafting | Transplanting | Bud | Anthesis |
|  | 17/03/19 | 22/03/19 | 09/10/19 | 19/11/19 | 21/06/21 | 06/07/21 |
| Days P | 0 | 5 | 201 | 41 | 580 | 15 |
| Days B | 0 | 5 | 206 | 247 | 827 | 842 |

*Note: “Days P” means days from the previous activity while “Days B” means days from the beginning of the investigation.*

The seed treatment which resulted in early germination and fast growth of rootstock coupled with matured scion and probably the good climate as well, led to the early flowering observed. With good agronomic practices and earlier grafting than we did, anthesis and fruiting could even be earlier than this observation. This result could encourage farmers to grow baobab.

At the time of anthesis, the height of the tree was only 170 cm as recorded in Table 2. The first branch was as low as 30 cm and the circumference of tree trunk at ground level amd at first branch 35.4 cm and 27.7 cm respective;y. These values are far lower than would be expected in nature. Tree stature would make management easy. The tree and flower are shown in Figs. 1 and 2.

**Table 2:** Data on tree at first anthesis

| Character | Measurement |
| --- | --- |
| The plant height at first flowering was | 170 cm |
| First branch height | 30 cm |
| Circumference at ground level | 35.4 cm |
| Circumference at first branch | 27.7 cm |
| Radius of the canopy | 141 cm |
| Graft union appearance | Smooth and not visible |

**Fig. 1:**
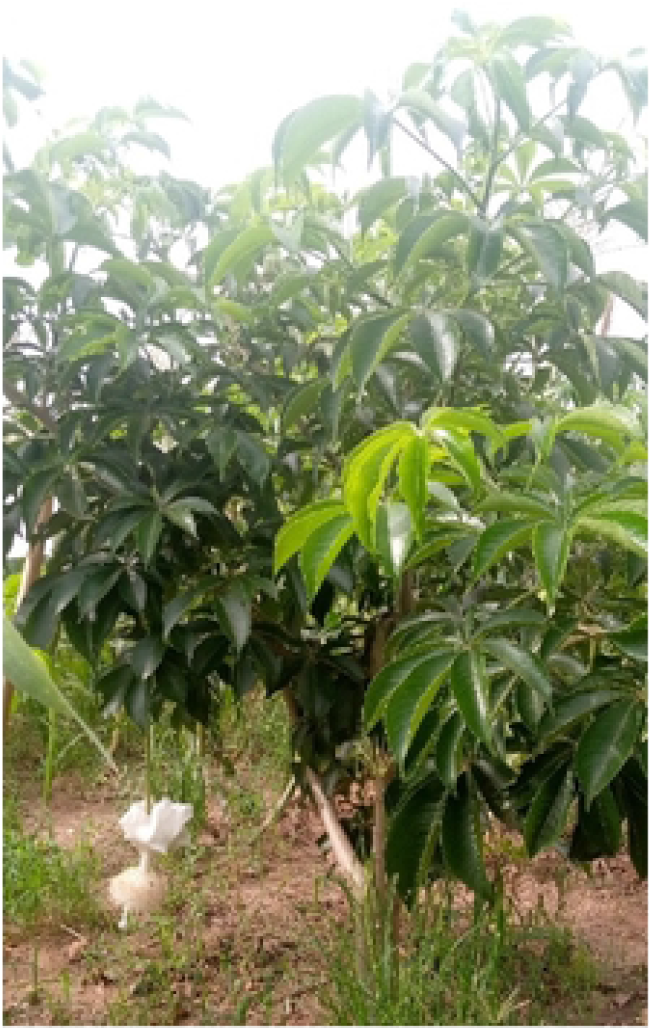
Flowering baobab tree.

**Fig. 2:**
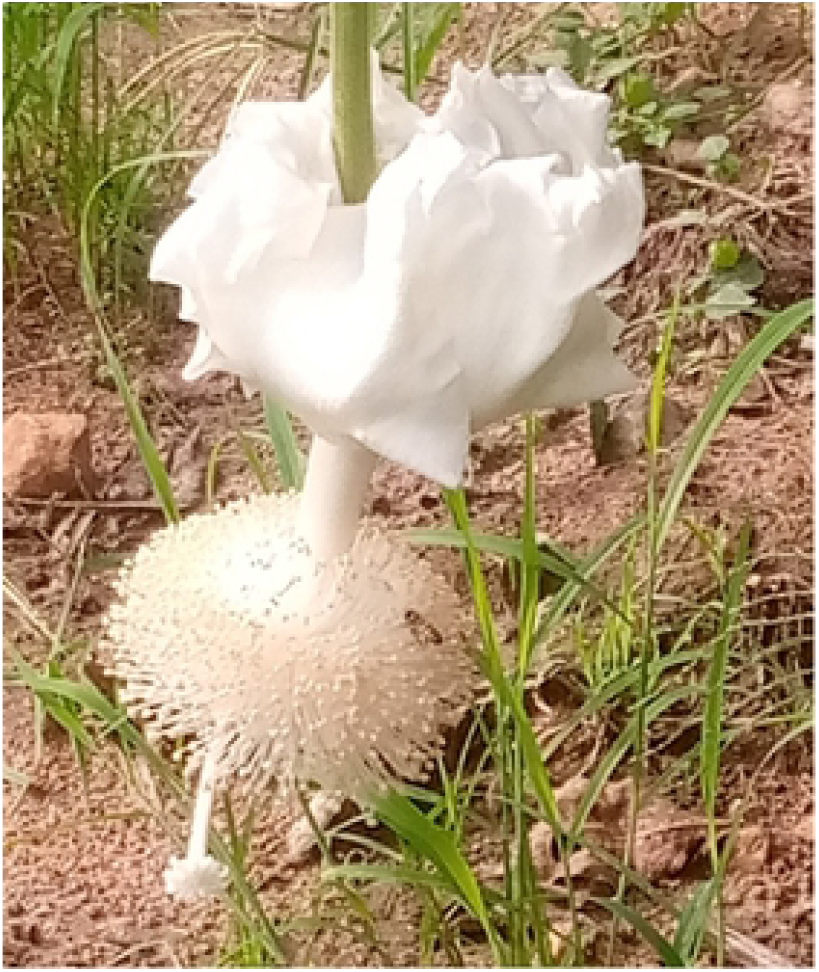
Baobab flower.

Although, flower was formed, it did not develop fruit. This is not unexpected because baobab is known to be self-incompatible (Venter *et al*, 2017). There is expectation of more flowers and fruiting would occur after few months.

## Conclusion

The studies ongoing has shown that there is great possibility of drastically reducing the maturity period of baobab to make it attractive to farmers to go into its cultivation. Although, cultural practices were not the best for our studies, anthesis occurred earlier than ever reported for baobab. Further investigations are therefore going on at Ho Technical University (HTU) to lead to its domestication.

## Acknowledgement

The authors appreciate the support given by the Vic-Chancellor of HTU (Professor Ben Q. Honyenuga) and the University management. We also thank both teaching and non-teaching staff of the Department of Agro Enterprise Development of HTU as well as students of the department who are working in one way or the other on the baobab project. We also thank the whole HTU community for celebrating our progress.

## Notes

### Competing Interest Statement

The authors have declared no competing interest.

